# Hierarchical transcriptional control regulates *Plasmodium falciparum* sexual differentiation

**DOI:** 10.1101/633222

**Authors:** Riëtte van Biljon, Roelof van Wyk, Heather J. Painter, Lindsey Orchard, Janette Reader, Jandeli Niemand, Manuel Llinás, Lyn-Marie Birkholtz

**Affiliations:** Department of Biochemistry, Genetics and Microbiology, Institute for Sustainable Malaria Control, University of Pretoria, Private Bag x20, Hatfield 0028, South Africa; Department of Biochemistry & Molecular Biology and the Huck Center for Malaria Research; Department of Chemistry, Pennsylvania State University, University Park, PA, 16802, USA

**Keywords:** Malaria, *Plasmodium*, gametocyte, gametocytogenesis, transcriptome, gene expression regulation, differentiation, sexual development

## Abstract

Malaria pathogenesis relies on sexual gametocyte forms of the malaria parasite to be transmitted between the infected human and the mosquito host but the molecular mechanisms controlling gametocytogenesis remains poorly understood. Here we provide a high-resolution transcriptome of *Plasmodium falciparum* as it commits to and develops through gametocytogenesis. The gametocyte-associated transcriptome is significantly different from that of the asexual parasites, with dynamic gene expression shifts characterizing early, intermediate and late-stage gametocyte development and results in differential timing for sex-specific transcripts. The striking transcriptional dynamics suggest strict transcriptional control during gametocytogenesis in *P. falciparum*, which we propose is mediated by putative regulators including epigenetic mechanisms (driving active repression of proliferation-associated processes) and a cascade-like expression of ApiAP2 transcription factors. The gametocyte transcriptome serves as the blueprint for sexual differentiation and will be a rich resource for future functional studies on this critical stage of *Plasmodium* development, as the intraerythrocytic transcriptome has been for our understanding of the asexual cycle.

## Introduction

Sustained malaria prevalence is ensured through continued human-to-mosquito transmission of *Plasmodium* parasites with *Plasmodium falciparum* being the causative agent of the most severe form of the disease in humans (1). The complex life cycle of *P. falciparum* encompasses development in the liver and erythrocytes of its human host and transmission by the female *Anopheline* mosquito. Two distinct developmental phases characterize intraerythrocytic development: rapid, cyclic asexual cell division manifesting in pathology, and the stochastic (<10%) sexual differentiation into gametocytes (2,3), which produces the non-replicative, mature, transmissible forms of the parasite. Whilst the intraerythrocytic developmental cycle (IDC) is relatively rapid (~48 h) and results in massive cell number expansion, sexual differentiation and development (gametocytogenesis) is a prolonged process (~10 days) in *P. falciparum* and is characterized by the development of the parasite through five morphologically distinct gametocyte stages (stages I-V) (4).

The processes of asexual replication and sexual differentiation in *Plasmodium* are associated with distinct patterns of gene expression that are tightly controlled through complex regulatory systems (5). These patterns have been investigated to some extent for asexual replication where *P. falciparum* parasites use both transcriptional (6–8) and post-transcriptional processes (9,10) to effect a cascade of coordinated, stage-specific gene expression (11,12). Despite the identification of some putative regulators of gene expression, including the Apicomplexan-specific ApiAP2 family of transcription factors (13–15) and epigenetic regulation of particular gene families (16,17), the specific mechanisms controlling transcriptional activation in the parasite are incompletely understood, with recent data clearly showing mRNA dynamics are also influenced by additional post-transcriptional mechanisms (18,19).

The mechanisms regulating commitment to gametocytogenesis have been somewhat clarified recently, with the discovery of host LysoPC restriction acting as an environmental factor driving gametocyte commitment (20). The AP2-G transcription factor acts as a molecular master switch of sexual commitment (21–24) and results in the expression of genes that drive entry into gametocytogenesis (22–26). The *ap2-g* gene is released from an epigenetically silenced state (27,28) through the antagonism of heterochromatin protein 1 (HP1) epigenetic silencing of the *ap2-g* locus by the gametocyte development protein 1 (GDV1). (29). Commitment to gametocytogenesis further requires stabilization of a subset of gametocyte-specific transcripts (18).

Despite these advances toward unravelling the mechanisms of commitment, the molecular functions governing subsequent gametocyte development and maturation remain poorly understood. Previously, deletion of certain ApiAP2 proteins has been shown to prevent progression of gametocyte development in the rodent parasite *P.* b*erghei* (30) and *P. falciparum* (31). Further, a subset of transcripts are translationally repressed by RNA binding proteins such as the Pumilio family protein (PUF2) during gametocytogenesis and ATP-dependent RNA helicase DDX6 (DOZI) and trailer hitch homolog (CITH) that repress female gametocyte transcripts needed to complete gametogenesis (32,33). However, systematic exploration of gene expression for *P. falciparum* gametocytogenesis has been limited to evaluation of the transcriptome (34–38) and proteome (35,39–41) at specific developmental timepoints. This includes the bifurcation in committing asexual parasites to gametocytogenesis (18,20,25), and evaluation of mature gametocytes in preparation for transmission (35,40,41). Current datasets that evaluate the complete gametocyte development process are sparse (36) and preclude dynamic evaluation of the transcriptomic profile associated with the extended gametocyte development process of *P. falciparum* parasites. Therefore, a time-resolved, high-resolution dataset capturing the transcriptome of each stage of gametocyte development would greatly enhance our ability to compare gene expression levels throughout the 10 days of gametocyte development and maturation.

Here we describe a comprehensive transcriptome analysis of *P. falciparum* parasites during all stages of sexual development at daily resolution. By measuring transcript abundance pre- and post-commitment, the transcriptional profile of gametocytes can be completely distinguished from that of asexual parasites. The data show marked shifts in transcript abundance associated with morphological stage transitions, indicating that gene expression occurs on a time scale consistent with developmental decisions underlying gametocyte development. We also show that post-commitment, the gametocyte transcriptome is shaped by specific epigenetic marks and ApiAP2 transcription factors. The gametocyte transcriptome provides a quantitative baseline of gene expression throughout sexual development and constitutes a highly valuable resource that could be exploited to further understand the molecular mechanisms governing sexual differentiation and maturation of the malaria parasite.

## Results

### Transition between sexual and asexual stages of development defined by transcriptome

*P. falciparum* NF54-*pfs16*-GFP-Luc (42) parasites were induced to form gametocytes after 1.5 cycles of asexual development (3 days) and monitored for the next 13 days to capture gametocyte commitment and development to mature stage V gametocytes (Fig 1A). Tight synchronization of the asexual parasites ensured coordinated gametocyte development, and gametocytes (stage I) were observed in culture from day 0 onwards (Fig 1B). Morphological evaluation showed a shift from a predominantly asexual parasite population to >60% gametocytes by day 3 of gametocytogenesis following the removal of asexual stages (Fig 1B).

**Fig 1:**
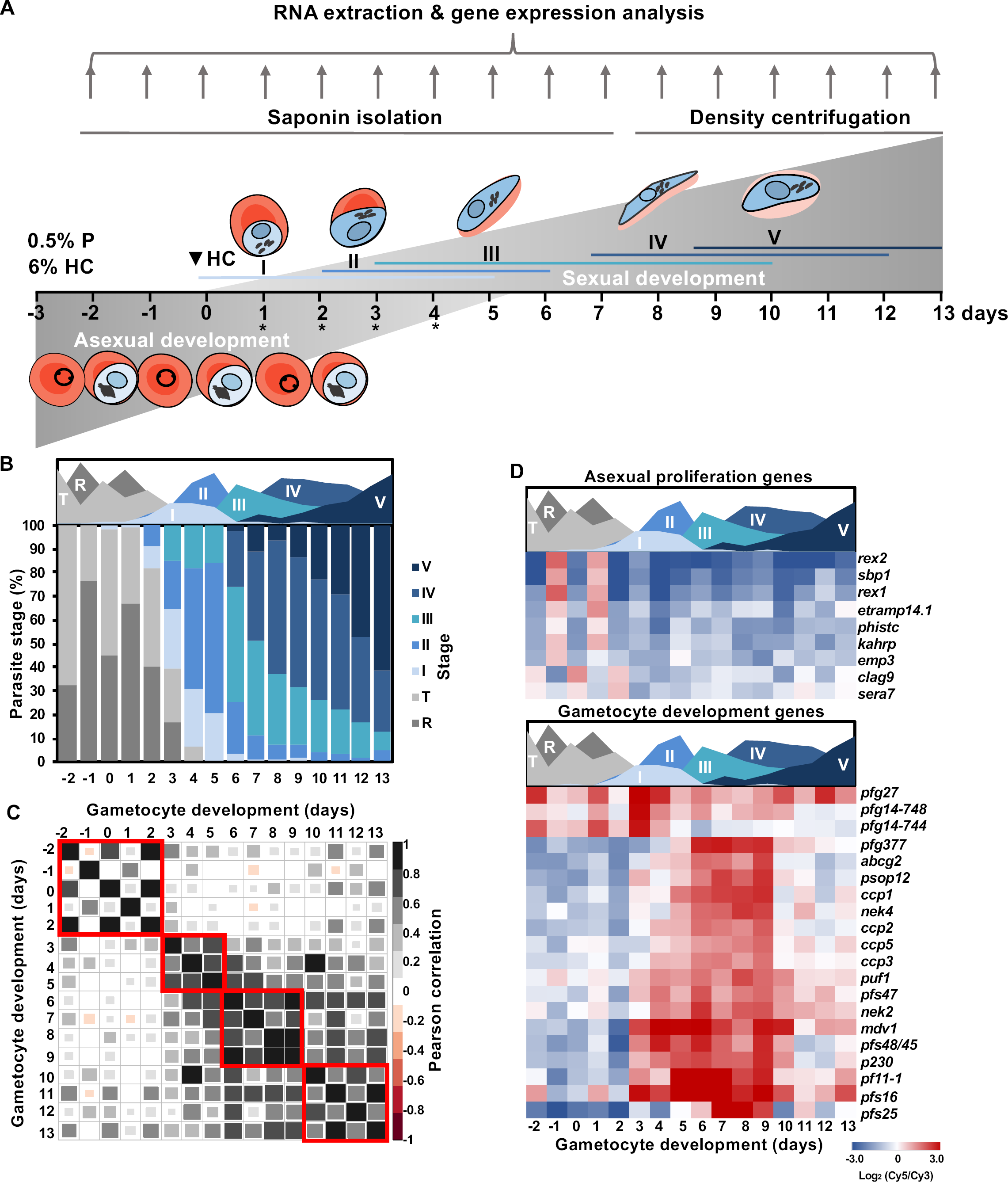
The developmental and associated transcriptomic profile of *P. falciparum* NF54 gametocytes from commitment to maturity. **(A)** Sampling and culturing strategy and stage distribution of parasites on each day of the time course. Colored lines indicate the presence of specific stages at different time points. Abbreviations indicate parasitemia (P) and hematocrit (HC) at induction, * indicate the addition of N-acetyl glucosamine (NAG) or 5% D-sorbitol. Parasite drawings were modified from freely available images (https://smart.servier.com/), under a Creative Commons Attribution 3.0 Unported Licence. **(B)** Morphological development was monitored from induction (day −2) over 16 days of development using Giemsa-stained thin-smear microscopy. The stage distribution for each day was calculated by counting ≥100 parasites on each day of monitoring. Legend: I-V indicates different stages of gametocyte development, R=ring and T=trophozoite stage asexual parasites. **(C)** Pearson correlation coefficients of the total transcriptomes obtained for each day of development. Red boxes indicate localized phases of increased correlation. **(D)** Expression of “gold standard” asexual and gametocyte genes (43) are shown for the gametocyte time course in heatmaps. **(A-D)** Area plot designates the timing of appearance and abundance of specific stages throughout the time course.

We measured mRNA abundance genome-wide using DNA microarrays at daily intervals, capturing expression values for 96-99% of the 5443 genes measured per timepoint (*P*<0.01, Supplementary File 1). Overall, the transcriptome of gametocytes is distinct from asexual parasites, as is evidenced by a clear shift in Pearson correlation between the transcriptomes of asexual parasites (day −2 to 2) and gametocytes (day 3 onward) (Fig 1C). Populations containing predominantly asexual parasites (days −2 to 2) were highly correlated across the first two 48 h cycles (r^2^=0.54-0.86, Supplementary File 2) and were characterized by periodic gene expression changes between the asexual ring and trophozoite stages (Fig 1C). From Day 3 onward, the transcriptional profiles diverged indicating a switch from asexual to sexual development, evidenced by a loss of the 48 h correlation pattern (Fig 1C). During subsequent days of gametocytogenesis, peak correlations were associated with developmental progression through stage I-II (days 3-5, r^2^=0.56-0.73), stage III-IV (days 6-9, r^2^=0.51-0.92), and mature stage V gametocytes (days 10-13, r^2^=0.50-0.84) (Fig 1C, Supplementary File 2).

Progression from asexual to sexual development was also clearly detectable in the expression profiles of individual genes required during asexual development (*e.g. kahrp (pf3d7_0202000)*) while sexual genes were only expressed during gametocyte development from Day 3 (43) (Fig 1D). The genes restricted to expression during sexual development include downstream targets of PfAP2-G (23) and markers associated with mature gametocyte sex-specificity (Fig 1D) (35) and 24 novel gametocyte-associated transcripts (Supplementary File 2). Among these transcripts were a putative ncRNA, three rRNAs and two tRNAs, suggesting that the expression of non-coding RNAs may not only play a role during gametocyte commitment (18) but also in gametocyte development and maturation in *P. falciparum*. Together, these data comprise a high-resolution *P. falciparum* blood stage developmental transcriptome that allows for the temporal evaluation of transcriptional abundance patterns associated with gametocyte commitment, development and maturation.

### The gametocyte-specific transcriptional program reflects the molecular landscape of gametocyte development

To associate temporal gene expression to particular events during gametocyte commitment and stage transitions throughout development, the full 16-day transcriptome dataset was K-means clustered revealing 2763 transcripts with overall decreased abundance (clusters 1-5) and 2425 with increased abundance during gametocytogenesis (clusters 6-10, Fig 2A). Therefore, gametocytogenesis relies on a more specialized program of gene expression compared to asexual development, with only 45% of transcripts showing increased abundance during gametocyte development (Fig 2A) compared to the 80-95% of transcripts increased during specific phases of asexual development (11,19,44). Interestingly, individual clusters showed specific patterns of gene expression throughout gametocyte development (Fig 2A), with transcript abundance during gametocytogenesis either decreased following asexual development (clusters 1-3, 1042 transcripts); maintained (clusters 4-5, 1721 transcripts) or increased (cluster 6-7, 1571 transcripts). Three clusters (clusters 8-10) show transcripts with specific peaks in expression evident during development, indicative of developmental regulation.

**Fig 2:**
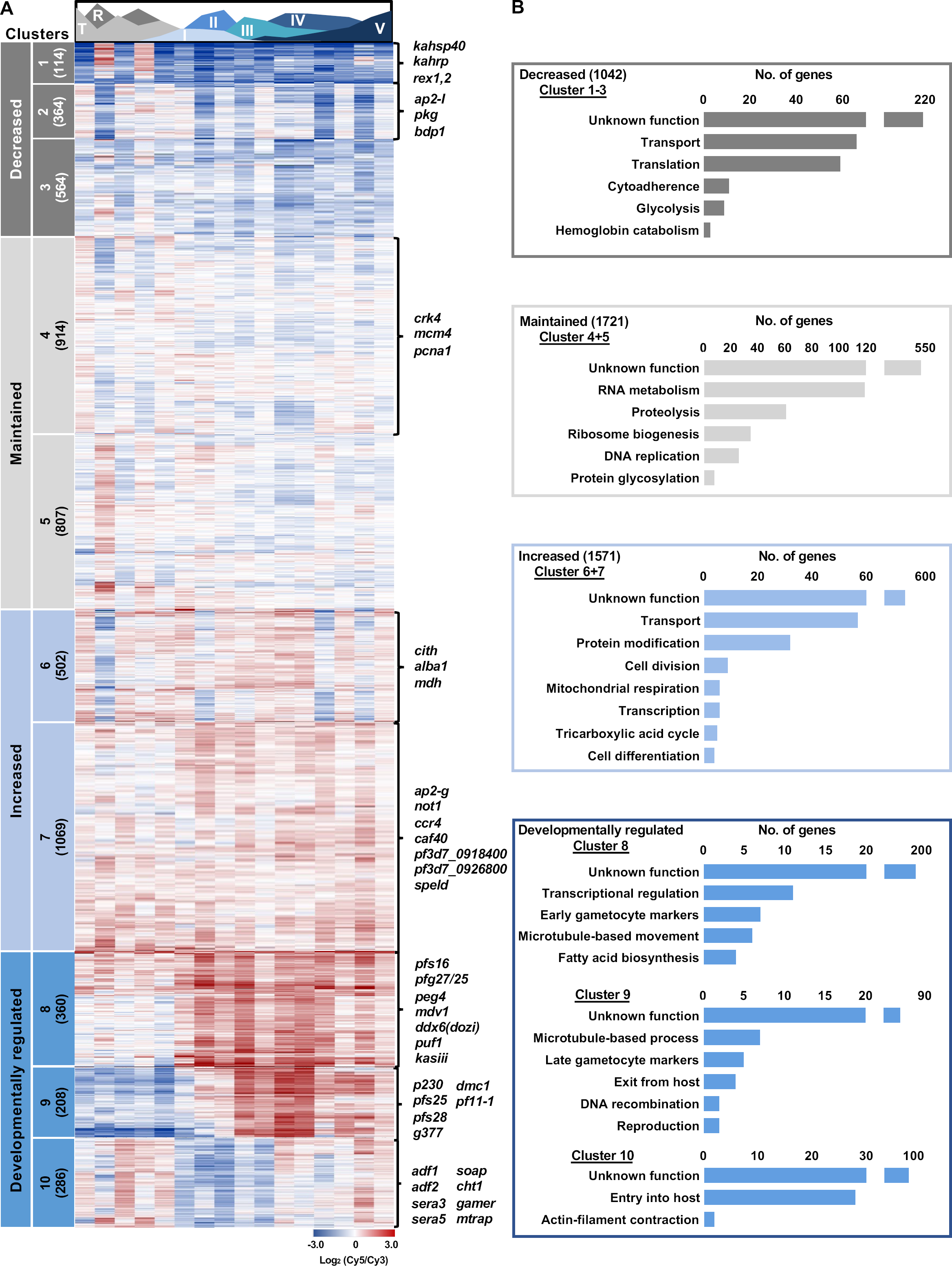
Distinct clusters of expression link to biological development of the *P. falciparum* gametocyte. Clusters of genes expressed during gametocyte development following K10 clustering of the total transcriptome. **(A)** The 10 clusters were grouped into phases of decreased, maintained, increased or developmentally regulated transcript abundance with number of transcripts per cluster indicated in brackets and genes of interest from specific clusters highlighted next to heatmaps. Area plot designates the timing of appearance and abundance of specific stages throughout the time course. **(B)** Biological processes of interest were selected from GO enrichment (Supplementary File 1) of each of the clusters (*P<*0.05) with the number of genes related to these functions shown for the groups of clusters in bar graphs with generic descriptions of the gene sets used to describe their function.

Cluster 1 is predominantly comprised of critical asexual stage transcripts which showed a decline in abundance during gametocytogenesis, with up to 5 log_2_ fold decreases in the expression of these transcripts between the ring and early gametocyte stages (Fig 2A). These transcripts include Maurer’s cleft proteins *e.g. rex1 (pf3d7_0935900)* and *rex2 (pf3d7_0936000)* as well as knob associated proteins that form part of the cytoadherence complex *(kahrp (pf3d7_0202000), kahsp40 (pf3d7_0201800))*, supporting earlier observations that the gametocytes mediate sequestration via different mechanisms than asexual parasites (45). Many of these cytoadherence associated transcripts are associated with heterochromatin protein 1 (HP1) occupancy during gametocyte development (46), and other HP1 (46,47) and H3K9me3 (17) repressed genes are also significantly enriched in cluster 1 (*P<*0.0001, Fisher’s exact test, genes listed in Supplementary File 3). This suggests asexual development-specific genes are actively repressed by epigenetic regulation throughout gametocyte development. Clusters 1-3 also contain transcripts involved in metabolic processes that are not critical to gametocyte development including genes encoding for enzymes of heme metabolism and glycolysis (Fig 2B, cluster 3, Supplementary File 1) as well as regulators of egress (*pkg (pf3d7_1436600)*) and invasion (*bdp1 (pf3d7_1033700)* and *ap2-i* (*pf3d7_1007700)*), all processes that are not necessary for gametocyte maturation (Fig 2A, cluster 2). Beyond these examples, clusters 1-3 also contain 214 unannotated genes that could be specifically required for asexual development only (Fig 2B).

Some transcripts show low abundance throughout gametocytogenesis (Fig 2A, clusters 4 and 5, average expression <0.1 log_2_(Cy5/Cy3), with amplitude change <0.5 log_2_(Cy5/Cy3)). These clusters include regulators of proliferation (*e.g.* origin of replication complex protein *mcm4 (pf3d7_1317100)*, proliferating cell antigen 1 (*pf3d7_1361900)* and cyclin dependent kinase *crk4 (pf3d7_0317200*)). By comparison, clusters with transcripts maintained at increased levels throughout commitment and development (Fig 2A, clusters 6 and 7, average log_2_(Cy5/Cy3) >0.31) included expected gene sets involved in the constitutive processes of macromolecular metabolism (*e.g.* DNA replication, protein modification and RNA metabolism Fig 2B, Supplementary File 1) (36,38). Interestingly, cluster 6 (and cluster 2) showed a high degree of cyclic oscillation in transcript abundance (Fig 2A). Many of these transcripts relate to transport, general cellular metabolism and homeostasis, functions in which fluctuation would not be unexpected (Fig 2B, Supplementary File 1). Importantly, cluster 7 also contained transcripts classified by gene ontology as involved in cellular differentiation *(caf40 (pf3d7_0507600), pf3d7_0918400, pf3d7_0926800* and *speld (pf3d7_1137800))* (GO:0030154, *P*=0.026, Fig 2B, S1 Table).

A significant proportion (15%) of the transcriptome associated with peak expression during specific stage-transitions in gametocyte development (Fig 2A, clusters 8-10), reminiscent of the phased expression typical of the asexual transcriptome (11,12). Transcripts involved in early-stage development increased from stage I-II in cluster 8 in a transcriptional profile often associated with targets of AP2-G (22,23,25). Transcripts in cluster 9 increased in abundance in the intermediate phase of development (stage III-IV) before the expression of transcripts required for development in mosquitoes in cluster 10 (stage V, *gamer* (*pf3d7_0805200*), *mtrap* (*pf3d7_1028700*), *cht1* (*pf3d7_1252200*), Fig 2A & B). The transcripts in clusters 8-10 are thus markers of biological transitions during gametocyte development. Clusters 6 & 8 are enriched for genes that contribute to the metabolic shift to mitochondrial metabolism (*e.g.* malate dehydrogenase *(mdh, pf3d7_0618500))* and fatty acid biosynthesis (*e.g.* β-ketoacyl-ACP synthase III (*kasIII, pf3d7_0618500))* (48,49) in gametocytes, followed by the emergence of processes related to cytoskeletal formation (clusters 8 & 9, Fig 2A &B, Table S1) that lead to the construction of a rigid subpellicular microtubule array during the sequestering stages (stages I-IV) of gametocytes (50). The microtubule array results in the characteristic crescent shape of the intermediate stages before the complex depolymerizes in stage V that is accompanied by the increased transcript abundance of actin depolymerization factors 1 and 2 (Fig 2A, cluster 10, *pf3d7_0503400, pf3d7_1361400*) to allow for a more deformable erythrocyte that can re-enter circulation (50). This cluster also includes the genes encoding the serine repeat antigens (*sera*) 3 and 5 *(pf3d7_0207800, pf3d7_0207600)* that play a role in egress in asexual parasites (51,52), implying that they may retain this role during gametocyte egress from the erythrocyte in the mosquito midgut. The striking temporal patterns of transcript abundance in clusters 8-10 suggests strict transcriptional regulation of these genes to ensure the timing of gametocyte sequestration, circulation and egress. Interestingly, these patterns are exhibited by parasites that need not fulfil any of these functions when grown *in vitro* in the absence of host-interactions, suggesting that transcription of these genes is hard-wired.

### Different regulatory modules enable sexual commitment and development

The time-resolved gametocyte transcriptome also allows interrogation of the expression of genes involved in sexual commitment throughout gametocyte development (18,20,25) (Fig 3). In total, previous reports produced a set of 1075 unique genes proposed to function as an “on switch” that characterizes gametocyte commitment (18,20,25). Of these, 680 genes (63%) also have increased transcript abundance during gametocyte development (Fig 3). These increased transcripts include those encoding epigenetic regulators involved in cell cycle control such as SIR2A (PF3D7_1328800) and SAP18 (PF3D7_0711400) that contribute to decreased DNA synthesis and a block in proliferation (53,54) necessary for the parasite to differentiate (Fig 3A). The remaining 395 transcripts are not increased in abundance during gametocyte development, suggesting that these transcripts are short lived and possibly only essential during gametocyte commitment. These short lived transcripts include *gdv1*, whose protein product prevents epigenetic repression of *ap2-g* during commitment (29), and *iswi* and *sn2fl*, which encode chromatin remodeling proteins (Fig 3A), that are expressed in sexually committed cells downstream of *ap2-g* (25). From our data we also identified a specific 5’ *cis*-regulatory motif (AGACA) that is enriched upstream of 539 genes with increased transcript abundance throughout gametocytogenesis (Fig 3B, Supplementary File 3), suggesting that transcriptional regulation of genes containing this motif could be important for gametocyte development. This AGACA motif has been associated with sexual commitment and development in previous datasets (18,55). In addition, a second motif (ATGTGTA) was also highly over-represented in the transcripts with increased abundance throughout gametocytogenesis (Fig 3B, S3 Table). This motif has been correlated with genes involved in DNA replication (56) and the significance of this enrichment in genes associated with differentiation is unclear and their *trans-*acting factors have not been identified (15,57).

**Fig 3:**
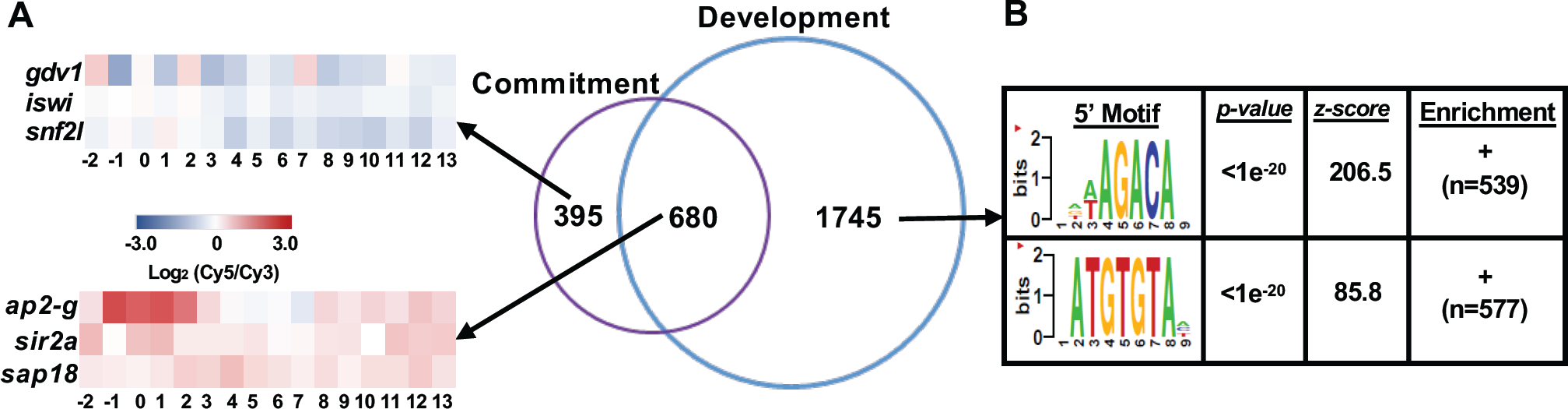
Commitment and development are distinctly regulated processes. **(A)** The genes increased in expression during commitment (18,20,25) were compared to transcripts increased in abundance during gametocytogenesis (Clusters 6-10, 2425 transcripts) with overlapping genes of interest: *sap18 (pf3d7_0711400), sir2a (pf3d7_1328800)* and genes only increased during commitment *gdv1 (pf3d7_0935400), iswi (pf3d7_0624600), sn2fl (pf3d7_1104200)* highlighted in heatmaps. **(B)** The 2429 genes expressed during gametocytogenesis also contained significantly enriched regulatory 5’ motifs identified using the FIRE algorithm (56).

### Transcriptional patterns characterize distinct transitions in gametocyte development

Apart from commitment to sexual development, the parasite also undergoes distinct developmental and transcriptional transitions during gametocyte development. The initial transition occurring in stage I gametocytes and regulating immature gametocyte development is characterised by increased transcript abundance in cluster 8 (Fig 4A), which showed a significant enrichment for genes involved in regulation of transcription (GO:0010468, 11 transcripts, *P=*0.029) including the specific ApiAP2 transcription factors *pf3d7_0404100, pf3d7_0516800, pf3d7_1429200* and the *myb1* transcription factor (*pf3d7_1315800)* (Fig 4A). Other genes with potential regulatory functions include a possible novel transcription factor, *pf3d7_0603600*, which contains an AT-rich interaction domain (IPR001606: ARID) and an uncharacterized RNA binding protein *(pf3d7_1241400)*. Proteins expressed by these two genes have been detected previously during gametocyte development (Fig 4A) (34,35,40,41). These proteins, along with the C-Myc binding protein MYCBP (PF3D7_0715100), are of interest for further study to determine their role in controlling gene expression during gametocyte development.

**Fig 4:**
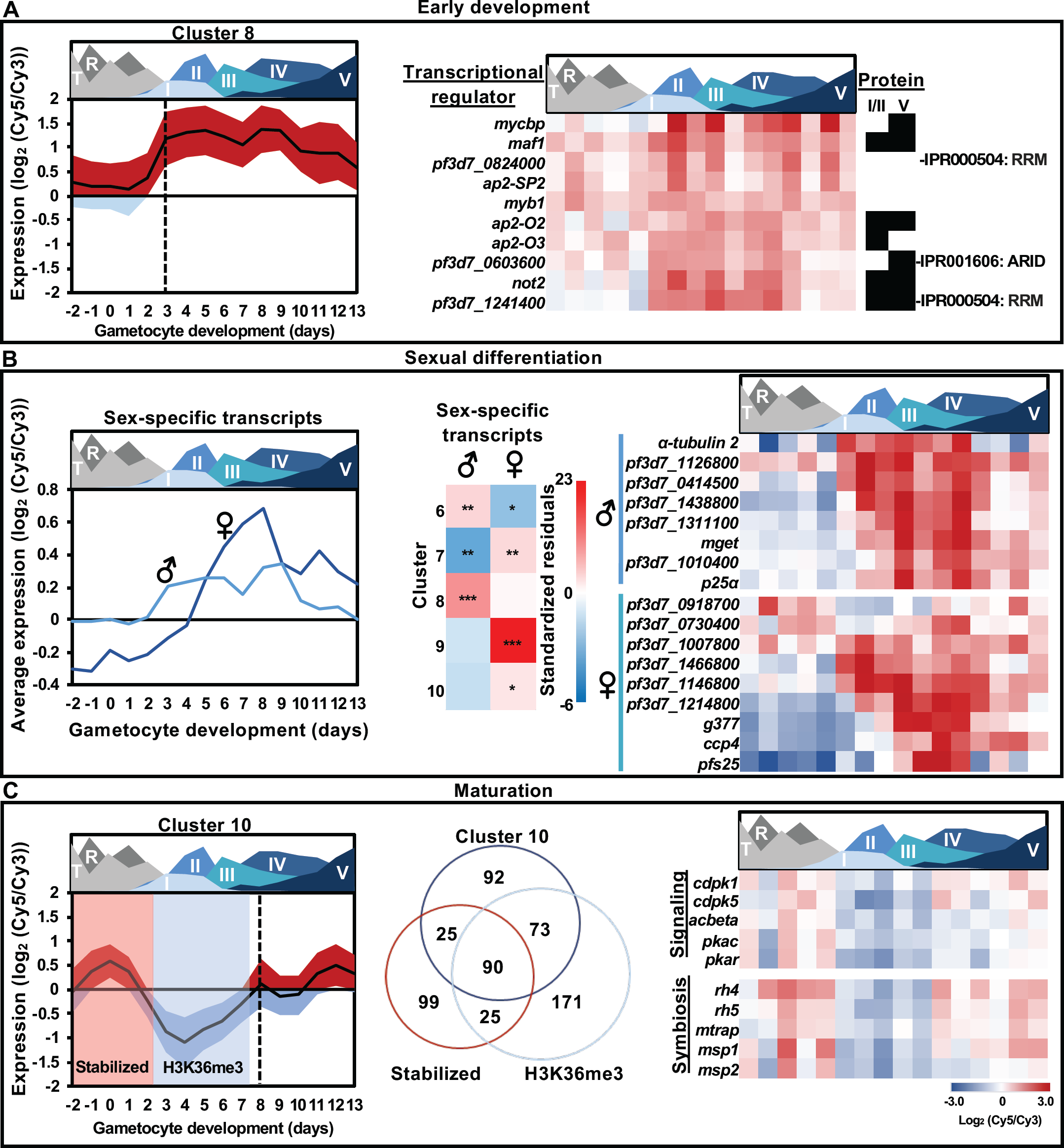
Stage-specific increases in gene expression contribute to the extended differentiation of *P. falciparum* gametocytes. **(A)** During stage I-III of development genes in cluster 8 sharply increased in expression (indicated with dotted line) with the abundance of these transcripts indicated by ribbon plot with mean±SD. GO enrichment of genes involved in regulation of transcription (GO:0010468, 11 transcripts, *P=*0.029) is present in this cluster, with presence of protein for these genes in stage I/II and V indicated in black (35,39–41) and the corresponding Interpro domains (https://www.ebi.ac.uk/interpro/) of proteins with unknown function indicated on the right. **(B)** The timing of sexually dimorphic transcript profiles (35) are shown in line graphs while the association of male-and female-enriched transcripts with specific clusters (6–10) are shown as standardized residuals and significance of these associations indicated (*P<*0.05*,0.001**,0.0001***, Fisher’s exact test). Genes of interest for each sex are highlighted in heatmaps next to male and female symbols. **(C)** The genes expressed during maturation (cluster 10) showed a significant association (*P<*0.0001, two-tailed Fisher’s exact test) with genes stabilized post-transcriptionally during commitment (18) and H3K36me3-associated genes in asexual development (16,73) before a sharp increase at stage IV-V of development (dashed line). Blocks indicate the timing of stabilization of the transcripts (18) or abundance of the H3K36me3 mark (65) and the overlap between the 3 datasets are indicated in the Venn diagram. Genes of interest within the three functional datasets are highlighted in heatmap. **(A-C)** Area plot designates the timing of appearance and abundance of specific stages throughout the time course.

A second outcome of the initial transition into gametocytogenesis is the determination of sex differentiation in *P. falciparum* parasites, which is proposed to be an PfAP2-G independent process that occurs at the very onset of commitment (18,35,40,41,58). However, sexually dimorphic gametocytes are only morphologically detectable by microscopy from stage III onwards (59). Our data indicate that the male-enriched transcripts from Lasonder *et al.* 2016 (35) show increased abundance earlier in development (stage I-II; 27% of cluster 8, *P<*0.0001, two-tailed Fisher’s exact test, Fig 4B, Supplementary File 3) compared to female transcripts. This observation is particularly interesting given the low ratio of male to female gametocytes in mixed populations (60), which suggests the rapid increase in male-enriched transcript abundance may be important for promoting male gametocyte development. These 98 male-enriched transcripts include 3 genes containing putative RNA binding motifs (*pf3d7_1126800, pf3d7_0414500, pf3d7_1438800*) that are highly abundant from stage I onwards, suggesting that these transcripts may be good biomarkers of early male differentiation as an alternative to α-tubulin II, which is expressed promiscuously in early gametocyte populations (61). Other male transcripts code for a meiotic protein (PF3D7_1311100), MGET (PF3D7_1469900), PF3D7_1010400 and P25? (PF3D7_1236600), which increase in abundance following the initial transcripts.

Female-enriched transcripts (35) peak in abundance only after sexual dimorphism is clearly discernible, from stage II-III onwards (Fig 4B) and are significantly overrepresented in the intermediate development cluster 9 (Fig 4B, Supplementary File 3, 76% of the cluster, *P<*0.0001, two-tailed Fisher’s exact test). Overall, this trend held true for the 158 female-enriched transcripts in cluster 9, including those encoding canonical female markers, *e.g.* osmiophilic body protein *g377* (*pf3d7_1250100)* (62,63), late-stage antigen *pfs25* (*pf3d7_1031000)* (35,63) and *ccp1-3 (pf3d7_1475500, pf3d7_1455800, pf3d7_1407000)* (35,63) and *ccp4 (pf3d7_0903800)* that was recently used to reliably type male and female gametocytes in late-stage gametocytes (64). We also detect a small subset of female-enriched transcripts (*pf3d7_0918700, imp2 (pf3d7_0730400), pf3d7_1007800, pf3d7_1466800, pf3d7_1146800, obc13 (pf3d7_1214800))* that are expressed earlier in gametocyte development (Fig 4B) and could potentially be important for female development before morphological differences are apparent.

The second transcriptional transition we observed coincides with the onset of gametocyte maturation from stage IV to V (Fig 4C). These transcripts show increased abundance in sexually committed asexual parasites as well as mature stage V gametocytes but have diminished abundance during the early and intermediate stages of gametocytogenesis (cluster 10, Fig 4C). This cluster was significantly enriched for transcripts stabilized during commitment (47% of transcripts, *P*<0.0001, two-tailed Fisher’s exact test) (18), as well as genes marked with H3K36me3 in asexual parasites (49% *P*<0.0001, Fisher’s exact test) (16). Interestingly, the epigenetic H3K36me3 mark is abundant during the intermediate stages of gametocyte development (65) and genes overlapping in the three datasets encode transcripts associated with the intracellular signalling machinery of the parasite (*cdpk1 (pf3d7_0217500), cdpk5 (pf3d7_1337800)* and adenylyl cyclase beta *(pf3d7_ 0802600*, (66)), along with cAMP-dependent protein kinase A catalytic and regulatory subunits (*pkac (pf3d7_0934800), pkar (pf3d7_1223100)* (Fig 4C). Of these, CDPK1 has been confirmed to function in de-repressing female gametocyte transcripts during parasite development in mosquitoes (67). Several of the genes in cluster 10 also have roles in invasion including the merozoite proteins *msp1, pf3d7_0930300, msp2, pf3d7_0206800, rh4, pf3d7_0424200, and rh5, pf3d7_0424100*, suggesting that invasion genes need to again be expressed for transition to gametogenesis in the mosquito. Overall, the gametocyte transcriptome reveals three major stages in gametocyte development (differentiation (Fig 4A), intermediate development (Fig 4B), maturation (Fig 4C)) that promote gametocyte development of *P. falciparum* parasites.

### ApiAP2 transcription factors are expressed at specific intervals during gametocytogenesis

To investigate the possible contribution of factors associated with transcriptional regulation to the observed stage-progressions during gametocytogenesis, we interrogated the expression of the genes encoding the ApiAP2 transcription factor family (Fig 5). Of the 27 family members, 15 genes encoding ApiAP2 transcription factors increased in transcript abundance during gametocyte development. Transcript abundance for *pf3d7_0404100, pf3d7_1350900, pf3d7_1449500, pf3d7_0802100, pf3d7_1429200* increased consistently throughout the time course (Figure S1). However, most ApiAP2-encoding transcripts increased in abundance at discrete intervals (Fig 5A) throughout gametocytogenesis. As expected, *ap2-g (pf3d7_1222600)* transcript abundance increased before the appearance of gametocytes (days −1 to 2). The target genes bound by AP2-G (23), peaked in transcript abundance directly following AP2-G peak abundance as expected, coinciding with stage I of gametocyte development (Supplementary Fig S2). Thereafter, three transcription factors *pf3d7_1408200, pf3d7_1317200* and *pf3d7_0611200* were increased during stage I to III of development (days 2-6). In the rodent-infectious malaria parasites *P. berghei* and *P. yoelii*, orthologs of the first two genes have been associated with gametocyte development through knockout studies (21,30). Three ApiAP2-encoding transcripts for *pf3d7_0516800, pf3d7_1222400, pf3d7_0934400* were increased in abundance from stage I to V of development (Fig 5A), following a pattern similar to the increased abundance of cluster 8 (Fig 4A). During the later stages, *pf3d7_1143100, pf3d7_1239200* and *pf3d7_0613800* were increased in abundance. Expression of PF3d7_1143100 is translationally repressed in *P. berghei* gametocytes (32), indicating that these transcription factors may not contribute to gene expression in *P. falciparum* gametocytes, but may instead have functional significance in subsequent development in the mosquito.

**Fig 5:**
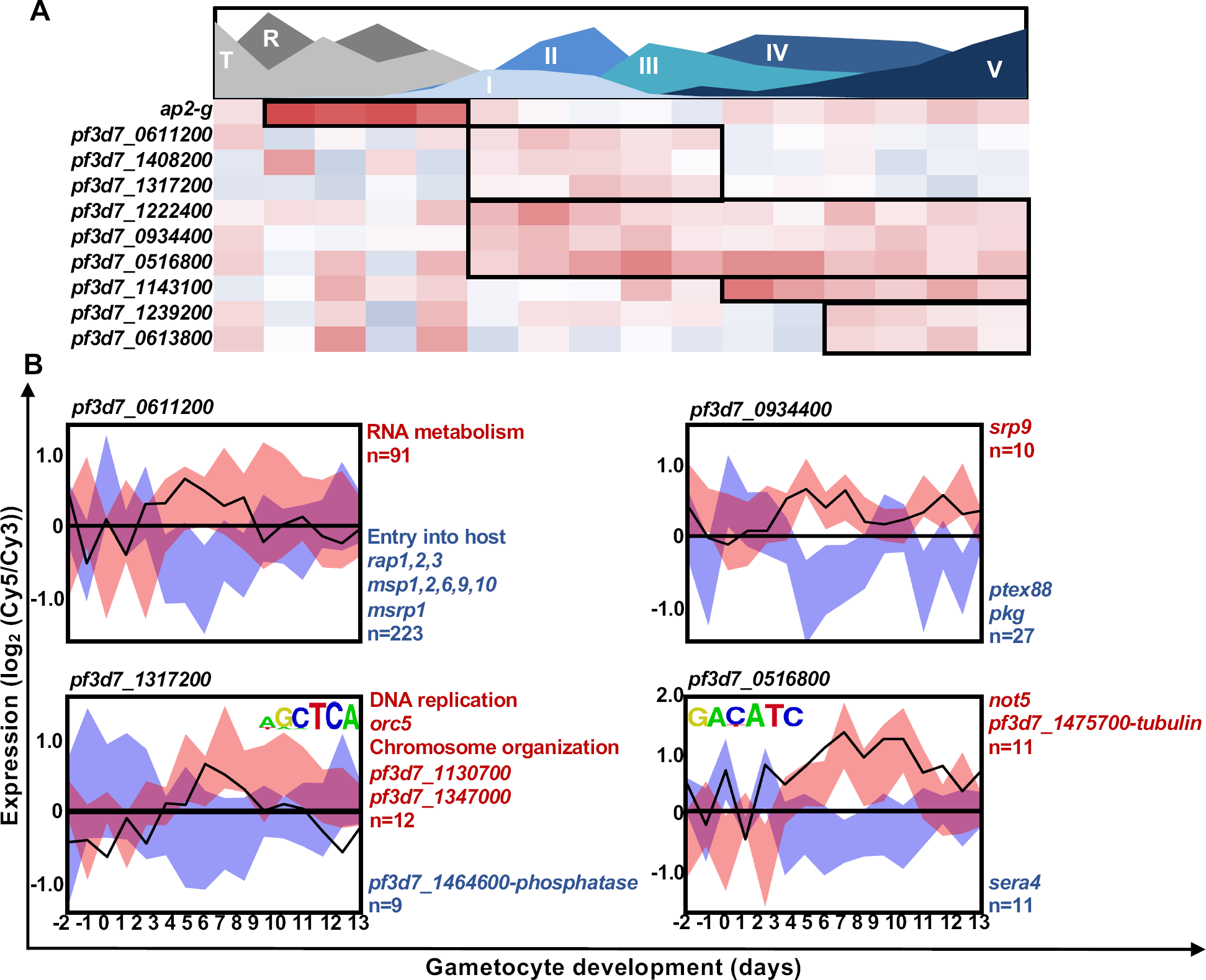
ApiAP2 transcription factors act as regulatory elements during gametocytogenesis. **(A)** ApiAP2 transcription factors increased in transcript abundance during gametocytogenesis were evaluated for their expression throughout gametocyte development with blocks indicating periods of increased abundance. Area plot designates the timing of appearance and abundance of specific stages throughout the time course. **(B)** The transcription factors were also probed for regulatory activity using coexpression analysis by GRENITS. Transcription factors with known binding sites (13), were probed against genes containing the transcription factor binding sites indicated or the total transcriptome if their binding site was unknown. The targets of each transcription factor are shown by shaded ribbons, with correlated transcripts indicated in red and anticorrelated transcripts indicated in blue. Generic functional terms describing enriched gene ontology terms or individual gene products are indicated in red (increased transcripts) or blue (decreased transcripts).

To associate functional regulation of gene sets due to the cascade-like increased abundance of the transcripts encoding the ApiAP2 transcription factors, the gametocyte transcriptome was analysed using Gene Regulatory Network Inference Using Time Series (GRENITS (68)) (Supplementary File 1) with the strongest predicted regulators shown in Fig 5B. From this analysis, *pf3d7_0611200*, which increased in abundance directly following *ap2-g*, coexpressed with 314 genes (probability linkage >0.6), 223 of which were anti-correlated for expression and functionally enriched for genes involved in host invasion (GO:0044409: entry into host, *P=*2.97e^−12^; Fig 5B). The large proportion of genes putatively repressed by this transcription factor would point to *pf3d7_0611200* acting as a repressive ApiAP2 transcription factor during gametocyte development, either alongside or instead of the *P. falciparum* ortholog of *pbap2-g2, pf3d7_1408200*. The second *apiap2* transcript increased in abundance, is *pf3d7_1317200*, the *P. falciparum* ortholog of *pbap2-g3*, coexpressed with 21 genes involved in cell cycle processes, including DNA replication (GO:0044786, *P=*0.0061) and chromosome organization (GO:0051276 *P=*0.0046). This ApiAP2 transcription factor has been previously associated with gametocyte development (31,69) and could be an important driver of stage II-III development. The two final ApiAP2 transcription factors are increased between stage I-V of development, with the first, *pf3d7_0934400*, showing mostly negative co-expression with its target genes (27/37 transcripts, including *pkg* and *ptex88 (pf3d7_1105600)*), suggesting this ApiAP2 transcription factor might also act as repressor. Secondly, the transcript of *ap2-o2* is increased in abundance throughout development but peaks at stage IV (day 8-9) of development and was predicted to regulate 22 target genes. Taken together, this data supports the involvement of successive expression of ApiAP2 transcription factors in a regulatory cascade during gametocyte development, as has been proposed for *P. berghei* gametocytes (21) and shows that this subsequent expression co-occurs with stage transition during *P. falciparum* gametocytogenesis.

## Discussion

We describe a high-resolution gametocyte transcriptome of malaria parasite differentiation from the asexual form through sexual commitment and all stages of development to mature stage V gametocytes. We find that gametocytogenesis in *P. falciparum* is a well-controlled process involving successive activation of regulatory processes that mediate development during stage-transition, ultimately resulting in a parasite poised for transmission. These observations emphasize that stage-specific gene expression is an essential feature of regulation of gene expression in *Plasmodium* spp. and is particularly true for the extended and morphologically diverse gametocyte development of *P. falciparum* parasites. The dynamic evaluation of the transcriptome allows for the construction of a more complete molecular roadmap for gametocyte development (Fig 6).

**Fig 6:**
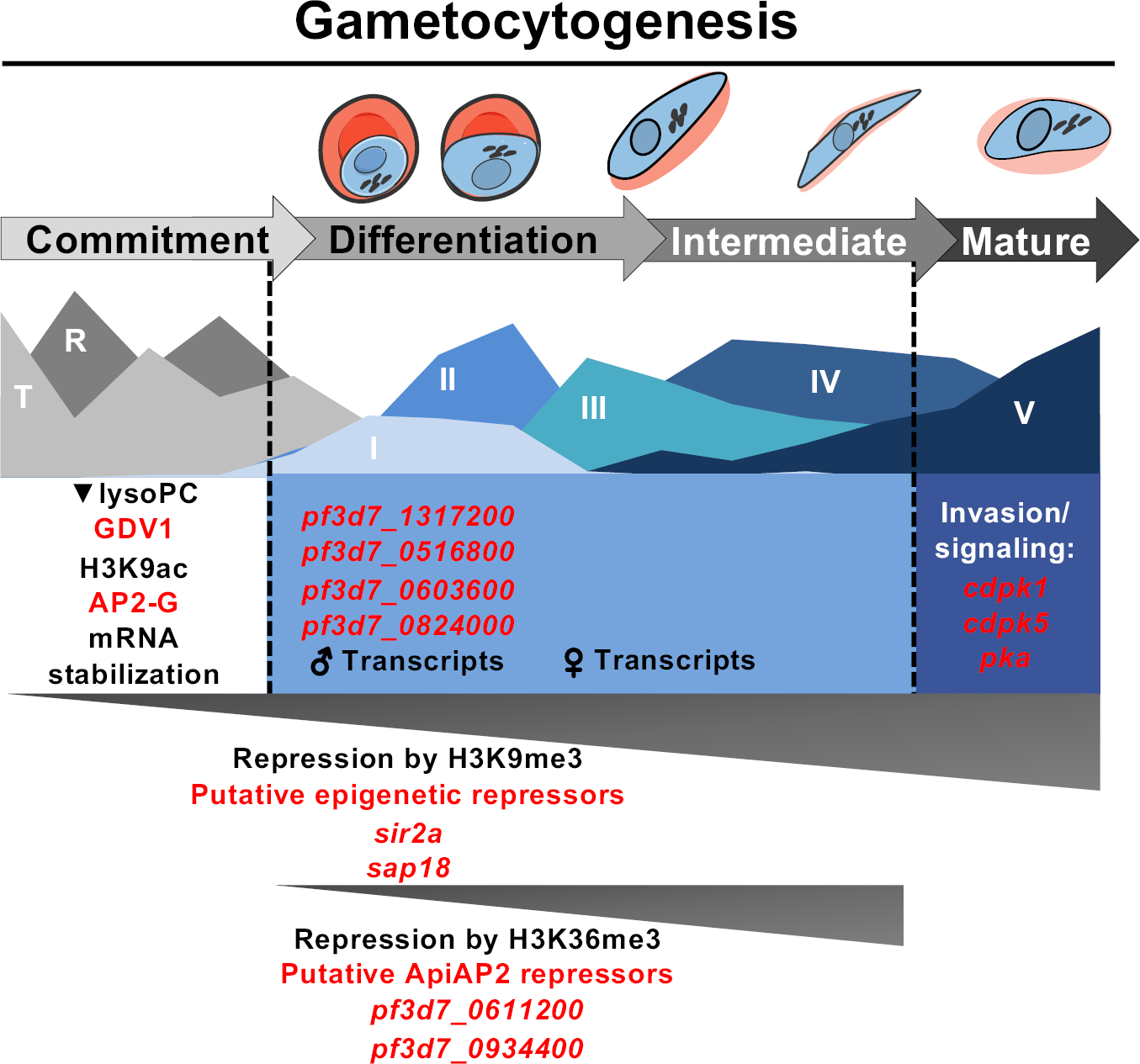
Molecular model of regulatory modules that shape cellular differentiation during gametocytogenesis. Specific regulatory events are mapped out over the extended gametocyte development of *P. falciparum* parasites. Molecular regulators are highlighted in red while specific events or epigenetic marks are shown in black. Colored blocks indicate the span of specific phases of transcript abundance, with dotted lines indicating transition points in gametocyte development and grey triangles indicate the timing of repressive mechanisms in gametocyte development. Parasite drawings were modified from freely available images (https://smart.servier.com/), under a Creative Commons Attribution 3.0 Unported Licence.

We propose that multiple transition points are passed during sexual differentiation and that mechanisms independent of initial sexual commitment during asexual proliferation are needed to ultimately result in completion of gametocyte development. First, committed parasites pass an initial transition point, whereby processes are initiated to drive early gametocyte development. This includes repression of genes typically associated with proliferation through epigenetic mechanisms (H3K9me3, HP1 occupancy), post-transcriptional regulation (18) and the activity of transcription factors that repress asexual-specific transcription (Fig 6). This transition point is also characterized by the increased transcript abundance of genes required during early and intermediate gametocyte development. A portion of these transcripts are expressed specifically in either sex, with an apparent delay between the peak abundance of male-specific and female-specific transcripts (Fig 6). It is possible that tracing the transcriptional dynamics within each sex separately would result in higher resolution data for this observation and resolve whether this is a true delay or underplayed by more complicated transcriptional dynamics that impact the RNA biology of the disparate sexes. As the gametocyte then reaches the critical transition for its pathology, the point of gametocyte maturation, a different set of genes increase in abundance. These genes are active in important processes specific to maturation into stage V gametocytes, including the switch from sequestration in the bone marrow to re-entering circulation and readying for transmission to the mosquito by involving the parasite’s intracellular signalling machinery.

A particularly interesting observation is the decreased abundance of important regulators of commitment, *ap2-g* and *gdv1*, as the parasite enters the early gametocyte stages (Fig 6). It is possible that the limited activity of these regulators might be essential for gametocytogenesis to occur normally, to allow the distinct patterns of gene expression we see here. It would be of interest to test what the effect of overexpression of one or both of these factors would be on gametocyte development. We also add to data on the transcriptional regulation in *P. falciparum* by the ApiAP2 transcription factor family downstream of AP2-G, affirming the presence of a transcription factor cascade enabling passage through gametocytogenesis as postulated for *P. berghei* (21). The involvement of *ap2-g2* and *pf3d7_0611200* in repressing transcription of asexual genes during gametocyte development (Fig 6) is also of particular interest for investigation, bringing into question if one or both of these factors fulfil this role in *P. falciparum* gametocytes. However, the possibility of novel regulators of transcription in early gametocyte development cannot be overlooked, with RNA binding proteins and the possible ARID transcription factor (Fig 6) good candidates for functional characterization.

The high-resolution transcriptome profile of *P. falciparum* gametocytes offers a complete molecular landscape of parasite differentiation. We identify putative regulators of mRNA dynamics facilitating a well-timed transcriptional program that prepares the parasite for transmission. The profile provides molecular identity to differences and similarities in asexual and sexual development that can be exploitable for pharmaceutical intervention. Finally, the stage-specific events that complicate transmission-blocking drug discovery are highlighted, 1) the immediate divergence of the gametocyte’s molecular profile from asexual development, 2) the later sexual dimorphism in intermediate stage development and 3) the apparent transcriptional divergence between immature and mature gametocytes. The gametocyte transcriptome further provides a valuable resource for further interrogation of the function of gene products and regulatory mechanisms important for gametocytogenesis in *P. falciparum*.

## Material and Methods

### Parasite culturing and sampling

*In vitro* cultivation of intraerythrocytic *P. falciparum* parasites and volunteer blood donation for human erythrocytes holds ethics approval from the University of Pretoria (EC120821-077). Asexual *P. falciparum* NF54 parasite cultures (NF54-*pfs16*-GFP-Luc,(42)) were maintained at 5-8% parasitemia 37°C in human erythrocytes at 5% hematocrit in RPMI 1640 medium supplemented with 25 mM HEPES, 0.2 % D-glucose, 200 μM hypoxanthine, 0.2% sodium bicarbonate, 24 μg/ml gentamicin with 0.5 % AlbuMAX^®^ II and incubated under hypoxic conditions (90% N_2_, 5% O_2_, and 5% CO_2_)(70). Synchronous asexual cultures (>95% synchronized ring-stage parasites) were obtained by three consecutive cycles of treatment with 5% D-sorbitol, each 6-8 h apart.

Gametocytogenesis was induced by employing a strategy of concurrent nutrient starvation and a decrease of hematocrit(70). Ring-stage parasite cultures were adjusted to a 0.5% parasitemia, 6% hematocrit in RPMI 1640 medium prepared as for growth of asexual parasites without additional glucose supplementation (day −3) and maintained under the same hypoxic conditions at 37°C without shaking. After 72 h, the hematocrit was adjusted to 3% (day 0). After a further 24 h, induction medium was replaced with medium containing 0.2% (w/v) D-glucose as the asexual parasites were removed daily with 5% D-sorbitol treatment for 15 minutes at 37°C and/or *N*-acetylglucosamine included in the culture medium for duration of the sampling.

All cultures were maintained with daily medium changes and monitored with Giemsa-stained thin smear microscopy and parasite stage distribution determined by counting ≥ 100 parasites per day. Parasite samples (30 ml of 2-3% gametocytemia, 4-6% hematocrit) were harvested daily for microarray analysis on days −2 to 13 following gametocyte induction. The samples harvested on days −2 to 7 were isolated from uninfected erythrocytes via 0.01% w/v saponin treatment for 3 minutes at 22°C while samples from day 8 to 13 were enriched for late stage gametocytes via density centrifugation using Nycoprep 1.077 cushions (Axis-Shield). Late stage gametocyte samples were centrifuged for 20 min at 800x*g* and the gametocyte containing bands collected(70). All parasite samples were washed with phosphate-buffered saline before storage at −80°C until RNA was isolated.

### RNA isolation, cDNA synthesis and microarray hybridization and scanning

Total RNA was isolated from each parasite pellet with a combination of TRIzol (Sigma Aldrich, USA) treatment and using a Qiagen RNeasy kit (Qiagen, Germany) as per manufacturer’s instructions. The quantity, purity and integrity of the RNA were evaluated by agarose gel electrophoresis and on a ND-2000 spectrophotometer (Thermo Scientific, USA). For each RNA sample, 3-12 µg total RNA was used to reverse transcribe and dye couple cDNA as described previously (71). The reference cDNA pool was constructed from a mixture of all the gametocyte samples used in the experiment in a 1:4 ratio with cDNA from a 6-hourly time course of asexual *P. falciparum* 3D7 parasites. For microarray hybridization, equal amounts of cDNA between 150 and 500 ng of Cy5 labeled sample and Cy3 labeled reference pool were prepared for hybridization as described previously (71). Arrays were scanned on an Agilent G2600D Microarray Scanner (Agilent Technologies, USA) with 5 μm resolution at wavelengths of 532 nm (Cy3) and 633 nm (Cy5). Linear lowess normalized signal intensities were extracted using the Agilent Feature Extractor Software version 11.5.1.1 using the GE2_1100_Jul11_no_spikein protocol and data was uploaded onto the Princeton University Microarray Database (https://puma.princeton.edu/).

## Data analysis

Signal intensities loaded on the Princeton University Microarray Database were filtered to remove background and unsatisfactory spots were flagged for removal using spot filters *P*<0.01 and log_2_ (Cy5/Cy3) expression values were used for further analysis. Euclidean distance clustered heatmaps were generated using TIGR MeV software version 4.9.0 (http://www.tm4.org/mev.html). The R statistical package (version 3.3.2) was used to calculate Pearson correlation coefficients and these were visualized using the Corrplot package. Data were divided into 10 clusters using K-means analysis following a within sum of squares test to determine the optimal number of clusters.

For functional analysis of genes, gene ontology enrichments were obtained for biological processes with *P<*0.05 using curated evidence using PlasmoDB Release v 33 (http://www.plasmodb.org/) and supplemented with annotation from MPMP (66) and Interpro (https://www.ebi.ac.uk/interpro/). Additional supplementary datasets for translationally repressed genes (33,35), transcripts involved in commitment (18,20,25) and gametocyte transcriptomes and proteomes (35,39–41) were probed for significant association with clusters of expression using a two-tailed Fisher’s exact test to calculate significant association between the datasets. For comparison between transcript abundance and histone post-translation modifications, supplementary information was sourced for ChIP-seq or ChIP-chip experiments from the Gene Expression Omnibus (GEO) datasets for H3K56ac, H4K5/8/12ac (72) as well as Salcedo-Amaya *et al.* for H3K9me3 (17), Jiang *et al.* 2014 for H3K36me3 (16) and Flueck *et al.* 2009 and Fraschka *et al.* for HP1 occupancy in *P. falciparum* parasites (46,47). The genes associated with the specific histone marks in each of the publications were then probed for association with specific clusters of expression using two-tailed Fisher’s exact tests and increased presence of the post-translational modification in gametocytes (65). To determine the involvement of ApiAP2 transcription factors in gametocyte development, the Gene Regulation Network Inference Using Time Series (GRENITS) package in R was applied (probability threshold >0.7) using the total transcriptome as possible regulated genes (68). The package uses Dynamic Bayesian Networks and Gibbs Variable Selection to construct a linear interaction model between gene expression profiles of putative “regulators” and “regulatees” over time-correlated data. Following the identification of 5 ApiAP2 transcription factors (*ap2-g* was not included in further predictive analysis) with putative regulatory activity, these transcription factors were re-probed as regulators, using genes containing the transcription factor’s binding site as possible regulated genes if the binding site had been determined (13). The number of links per model, per threshold was evaluated to determine the set probability threshold for the regulated genes of each transcription factor. The online FIRE algorithm (56) was used to identify enriched regulatory motifs in genes of interest.

## Data availability

The microarray data has been submitted to the GEO with accession number GSE104889 (www.ncbi.nlm.nih.gov/geo/).

## Author Contributions

RvB and LMB conceived the study. RvB, LO conducted experiments, HP contributed data, RvW, RvB, JR performed the analyses. RvB, RvW, JN, HP, ML and LMB interpreted results. RvB, ML, LMB wrote the paper with inputs from the other authors. All co-authors approved the final version of the paper.

## Acknowledgements

The UP ISMC acknowledges the South African Medical Research Council (SA MRC) as Collaborating Centre for Malaria Research. This work was supported by the South African Research Chairs Initiative of the Department of Science and Technology, administered through the South African National Research Foundation (UID 84627) and the European Commission “EviMalar” (no 242095) to LMB. We would like to acknowledge the Centre for Bioinformatics and Computational Biology (University of Pretoria) and the Centre for High Performance Computing South Africa for the use of servers in gene regulatory network construction.

## Supporting information captions

**Supplementary Fig S1. Transcript abundance of ApiAP2 transcription factors during *P. falciparum* gametocyte development**

**Supplementary Fig S2. Transcript abundance of *ap2-g* and downstream genes (identified in Josling *et al.* 2019)**

**Supplementary File 1. Total microarray data with GO enrichment pertaining to Figure 1&2**

**Supplementary File 2. Correlation of microarray time points and gametocyte markers pertaining to Figure 1**

**Supplementary File 3. Cross-dataset comparison and functional enrichment pertaining to Figures 2-5**

