## Supplemental figures for "Hierarchical transcriptional control regulates *Plasmodium falciparum* sexual differentiation"

### Slide 1
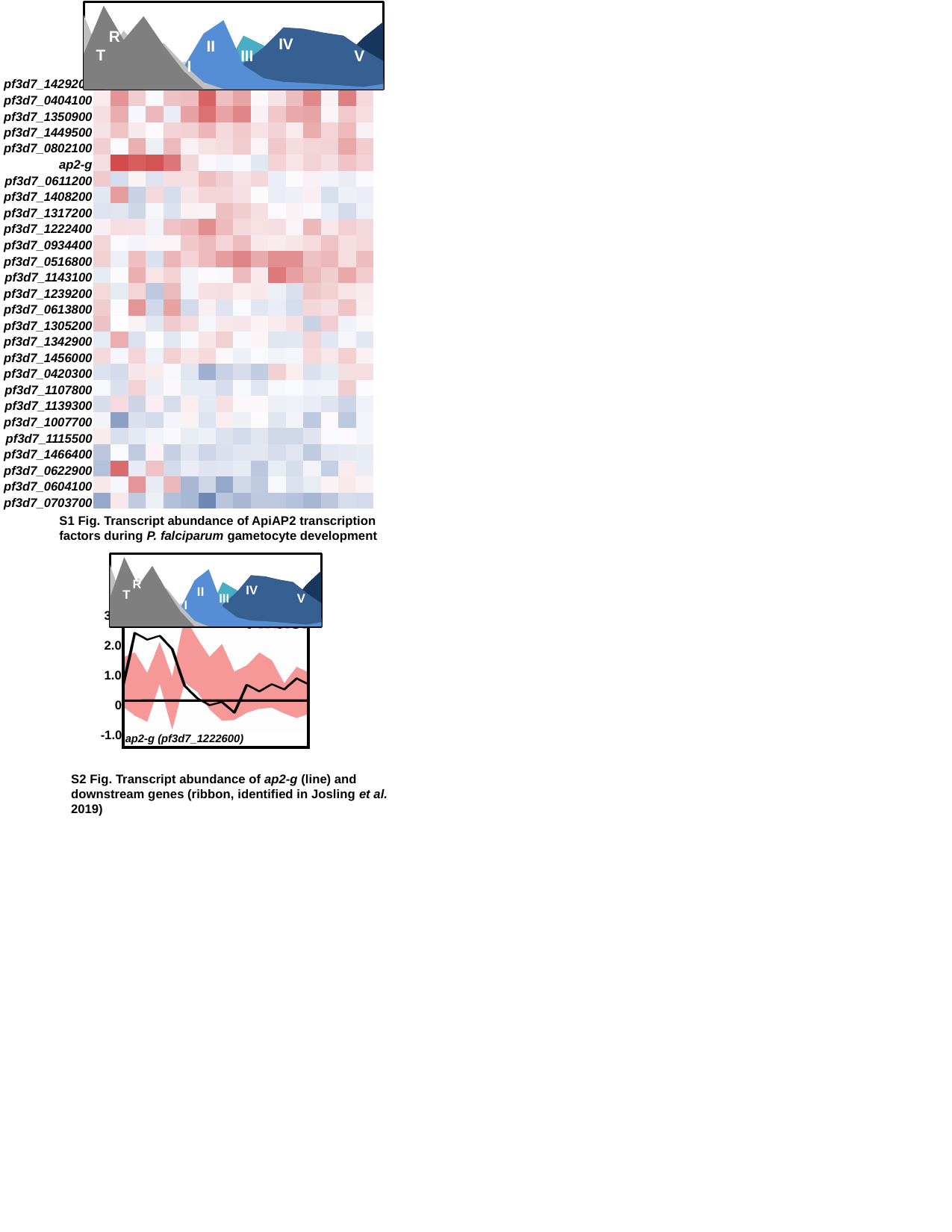

#### Chart
| Category | R | T | II | IV | III | V | I |
|---|---|---|---|---|---|---|---|
| -2 | 32.25806451612903 | 67.74193548387096 | 0.0 | 0.0 | 0.0 | 0.0 | 0.0 |
| -1 | 76.31578947368422 | 23.684210526315788 | 0.0 | 0.0 | 0.0 | 0.0 | 0.0 |
| 0 | 44.881889763779526 | 53.54330708661418 | 0.0 | 0.0 | 0.0 | 0.0 | 1.574803149606299 |
| 1 | 66.66666666666666 | 32.25806451612903 | 0.0 | 0.0 | 0.0 | 0.0 | 1.1235955056179776 |
| 2 | 40.0 | 42.22222222222222 | 11.39240506329114 | 0.0 | 3.79746835443038 | 0.0 | 9.0 |
| 3 | 16.455696202531644 | 22.78481012658228 | 20.253164556962027 | 0.0 | 15.18987341772152 | 0.0 | 25.31645569620253 |
| 4 | 0.0 | 6.153846153846154 | 50.76923076923077 | 0.0 | 18.461538461538463 | 0.0 | 24.615384615384617 |
| 5 | 0.0 | 0.0 | 63.0 | 0.0 | 16.0 | 0.0 | 21.0 |
| 6 | 0.0 | 0.0 | 21.875 | 23.958333333333336 | 48.95833333333333 | 2.083333333333333 | 3.125 |
| 7 | 0.0 | 0.0 | 10.0 | 38.0 | 40.0 | 11.0 | 1.0 |
| 8 | 0.0 | 0.0 | 6.481481481481481 | 56.481481481481474 | 29.629629629629626 | 6.481481481481481 | 0.9259259259259258 |
| 9 | 0.0 | 0.0 | 5.714285714285714 | 55.23809523809524 | 23.809523809523807 | 13.333333333333334 | 1.9047619047619049 |
| 10 | 0.0 | 0.0 | 4.49438202247191 | 51.68539325842697 | 21.34831460674157 | 22.47191011235955 | 0.0 |
| 11 | 0.0 | 0.0 | 3.1914893617021276 | 48.93617021276596 | 19.148936170212767 | 28.723404255319153 | 0.0 |
| 12 | 0.0 | 0.0 | 1.9607843137254901 | 36.27450980392157 | 14.705882352941178 | 47.05882352941176 | 0.0 |
| 13 | 0.0 | 0.0 | 4.651162790697675 | 25.581395348837212 | 8.13953488372093 | 61.627906976744185 | 0.0 |R
IV
II
T
III
V
I
| |
| --- |
| pf3d7\_1429200 |
| --- |
| pf3d7\_0404100 |
| pf3d7\_1350900 |
| pf3d7\_1449500 |
| pf3d7\_0802100 |
| ap2-g |
| pf3d7\_0611200 |
| pf3d7\_1408200 |
| pf3d7\_1317200 |
| pf3d7\_1222400 |
| pf3d7\_0934400 |
| pf3d7\_0516800 |
| pf3d7\_1143100 |
| pf3d7\_1239200 |
| pf3d7\_0613800 |
| pf3d7\_1305200 |
| pf3d7\_1342900 |
| pf3d7\_1456000 |
| pf3d7\_0420300 |
| pf3d7\_1107800 |
| pf3d7\_1139300 |
| pf3d7\_1007700 |
| pf3d7\_1115500 |
| pf3d7\_1466400 |
| pf3d7\_0622900 |
| pf3d7\_0604100 |
| pf3d7\_0703700 |
S1 Fig. Transcript abundance of ApiAP2 transcription factors during P. falciparum gametocyte development
#### Chart
| Category | R | T | II | IV | III | V | I |
|---|---|---|---|---|---|---|---|
| -2 | 32.25806451612903 | 67.74193548387096 | 0.0 | 0.0 | 0.0 | 0.0 | 0.0 |
| -1 | 76.31578947368422 | 23.684210526315788 | 0.0 | 0.0 | 0.0 | 0.0 | 0.0 |
| 0 | 44.881889763779526 | 53.54330708661418 | 0.0 | 0.0 | 0.0 | 0.0 | 1.574803149606299 |
| 1 | 66.66666666666666 | 32.25806451612903 | 0.0 | 0.0 | 0.0 | 0.0 | 1.1235955056179776 |
| 2 | 40.0 | 42.22222222222222 | 11.39240506329114 | 0.0 | 3.79746835443038 | 0.0 | 9.0 |
| 3 | 16.455696202531644 | 22.78481012658228 | 20.253164556962027 | 0.0 | 15.18987341772152 | 0.0 | 25.31645569620253 |
| 4 | 0.0 | 6.153846153846154 | 50.76923076923077 | 0.0 | 18.461538461538463 | 0.0 | 24.615384615384617 |
| 5 | 0.0 | 0.0 | 63.0 | 0.0 | 16.0 | 0.0 | 21.0 |
| 6 | 0.0 | 0.0 | 21.875 | 23.958333333333336 | 48.95833333333333 | 2.083333333333333 | 3.125 |
| 7 | 0.0 | 0.0 | 10.0 | 38.0 | 40.0 | 11.0 | 1.0 |
| 8 | 0.0 | 0.0 | 6.481481481481481 | 56.481481481481474 | 29.629629629629626 | 6.481481481481481 | 0.9259259259259258 |
| 9 | 0.0 | 0.0 | 5.714285714285714 | 55.23809523809524 | 23.809523809523807 | 13.333333333333334 | 1.9047619047619049 |
| 10 | 0.0 | 0.0 | 4.49438202247191 | 51.68539325842697 | 21.34831460674157 | 22.47191011235955 | 0.0 |
| 11 | 0.0 | 0.0 | 3.1914893617021276 | 48.93617021276596 | 19.148936170212767 | 28.723404255319153 | 0.0 |
| 12 | 0.0 | 0.0 | 1.9607843137254901 | 36.27450980392157 | 14.705882352941178 | 47.05882352941176 | 0.0 |
| 13 | 0.0 | 0.0 | 4.651162790697675 | 25.581395348837212 | 8.13953488372093 | 61.627906976744185 | 0.0 |R
IV
II
T
III
V
I
 3.0
 2.0
 1.0
 0
-1.0
ap2-g (pf3d7_1222600)
S2 Fig. Transcript abundance of ap2-g (line) and downstream genes (ribbon, identified in Josling et al. 2019)
